# Polyunsaturated fatty acids alter the formation of lipid droplets and eicosanoid production in *Leishmania* promatigotes

**DOI:** 10.1101/2022.07.01.498279

**Authors:** Yasmin Monara Ferreira de Sousa Andrade, Monara Viera de Castro, Victor de Souza Tavares, Rayane da Silva Oliveira Souza, Lúcia Helena Faccioli, Jonilson Berlink Lima, Carlos Arterio Sorgi, Valéria de Matos Borges, Théo Araújo-Santos

**Author notes:** Corresponding author: Théo Araújo-Santos, Ph.D., Address: Laboratório de Agentes Infecciosos e Vetores, Centro das Ciências Biológicas e da Saúde, Universidade Federal do Oeste da Bahia - Campus Reitor Edgard Santos, Rua da Prainha, 1326, Morada Nobre, Barreiras – Ba., zip code: 47810-047.

## Abstract

*Leishmania* parasites contain **l**ipid droplets (LD, or lipid bodies) and the molecular machinery responsible for synthesizing prostaglandins (PGs) and other bioactive lipids. We studied the effects of polyunsaturated fatty acids (PUFA) on LD biogenesis and eicosanoid production in distinct *Leishmania* species associated with different clinical forms of leishmaniasis. We also compared structural models of human-like cyclooxygenase-2 (GP63) and prostaglandin F synthase (PGFS) proteins of *Leishmania*, and we evaluated their enzymatic production in logarithmic and stationary growth phases of procyclic *L. amazonensis, L. braziliensis* and *L. infantum*. PUFAs modulate the formation of LDs in *L. braziliensis* and *L. infantum. Leishmania* species with equivalent clinical manifestations and tissue tropism had same protein mutations in GP63 and PGFS. No differences in GP63 production were observed among *L. amazonensis, L. braziliensis* and *L. infantum*, however increased PGFS production was detected during the parasite differentiation. Stimulation with arachidonic acid resulted in highly elevated production of hydroxyeicosatetraenoic acids compared to prostaglandins quantified by LC-MS/MS. The present findings open new perspectives on the role of eicosanoid metabolism in *Leishmania* and could contribute to the development of novel antiparasitic drugs.

## 1. Introduction

Lipid mediators are bioactive molecules derived from the metabolism of polyunsaturated fatty acids (PUFA) (Jordan and Werz, 2021). The most common lipid mediator precursors are derived from arachidonic (AA), eicosapentanoic (EPA), and docosahexaenoic (DHA) acids. (Jordan and Werz, 2021). Trypanosomatids, including *Leishmania*, can metabolize AA to eicosanoids by way of specific enzymes, such as cyclooxygenase (COX) and prostaglandin synthases (PG synthases) (Estrada-Figueroa et al., 2018; Kubata et al., 2007; Tavares et al., 2021). In addition, specialized lipid mediators are also identified in *Trypanosoma cruzi* (Colas, 2018; Paloque et al., 2019a; Tavares et al., 2021).

Lipid mediators are produced in the cytosol and in organelles termed lipid droplets (LD, lipid bodies) (Araújo-Santos et al., 2014; Bozza et al., 2011; de Almeida et al., 2018; Toledo et al., 2016), which are present in almost all organisms, including in trypanosomatid protozoa (Araújo-Santos et al., 2014; Olzmann and Carvalho, 2019; Onal et al., 2017; Tavares et al., 2021; Toledo et al., 2016). LDs are active sites for eicosanoid metabolism (Araújo-Santos et al., 2014; Bozza et al., 2011). Although protozoan parasites are known to produce a variety of specialized lipid mediators, studies investigating the role of these mediators in parasite biology and host-parasite interaction remain scarce.

Parasites possess the necessary machinery to synthesize lipid mediators (Kubata et al., 2007). *Leishmania* can metabolize AA to prostaglandins using PG synthases present in LDs (Araújo-Santos et al., 2014). Recently, the glycoprotein of 63 kDa (GP63) was described as a Cox-like enzyme responsible to convert AA to prostaglandin in *L. Mexicana* (Estrada-Figueroa et al., 2018). In addition to COX, trypanosomatids contain enzymes capable of synthesizing other eicosanoids, such as PGE_2_ and PGF_2α_. Both *T. cruzi* trypomastigotes and *L. infantum* respond to exogenous AA stimulation by producing prostaglandins (Araújo-Santos et al., 2014; Toledo et al., 2016). However, the presence of AA was not shown to alter PGF_2α_ synthase (PGFS) production in *L. infantum* (Araújo-Santos et al., 2014). *L. infantum* LDs are capable of synthesizing PGF2α, a mediator responsible for increasing parasite viability in the initial moments of infection via an as yet unknown mechanism (Araújo-Santos et al., 2014). *L. braziliensis* promastigotes and amastigotes also express PGFS, which may improve parasites fitness (Alves-Ferreira et al., 2020).

*Trypanosoma cruzi* trypomastigotes synthesize RvD1, RVE2 and RvD5, lipid mediators involved in the resolution of the inflammatory process (Colas, 2018). However, the role played by these mediators in the course of infection remains to be clarified. On the other hand, *T. cruzi* can synthesize and release thromboxane A_2_, which may exacerbate infection (Ashton et al., 2007). Although the presence of a thromboxane receptor has been demonstrated in *T. cruzi*, its function has not been determined (Mukherjee et al., 2014). Another pathway of lipid mediator production was recently described through the identification of proteins (CYP1, CYP2 and CYP3) similar to cytochrome P450 (CYP450) in the genome of *L. infantum*, which appear to be responsible for specialized lipid precursors in this parasite species (Paloque et al., 2019b).

Advances have been made in our understanding of the metabolism of bioactive lipids found in parasites, as well as mechanisms involving LDs. Leishmaniasis presents a diversity of clinical forms and symptoms related to specific *Leishmania* species. Herein we compared the formation of LDs and the production of eicosanoids in different New World *Leishmania* species using PUFA precursors as stimulant. Differences were identified in lipid metabolism among the *Leishmania* species investigated, and the enzymes related to eicosanoid production were described.

## 2. Methods

### 2.1 Parasites

*Leishmania infantum* (MCAN/BR/89/BA262) promastigotes were maintained for 7–9 days in hemoflagellate culture medium (HO-MEM) supplemented with 10% fetal bovine serum until reaching stationary phase (Araújo-Santos et al., 2014). For the cultivation of *L. amazonensis* (MHOM/BR/1987/BA125) and *L. braziliensis* (MHOM/BR/01/BA788), parasites were maintained in Schneider’s insect medium supplemented with 20% fetal bovine serum, L-glutamine, 20mM penicillin (100 U/ml) and streptomycin (0.1 mg/ml) at 26°C for 6 days until reaching stationary phase.

### 2.2 Stimulation of *Leishmania*

*Leishmania spp* in the third day of axenic cultures (logarithmic-phase promastigote) were used to perform the experiments. In 96 wells plates, 1×10^6^ parasites/well were either treated with progressively higher doses of EPA, DHA (3.75, 7.5, 15, 30µM) or AA (15µM), or with ethanol 0,3% v/v (vehicle), or medium alone (control), for 1h. Next, parasites were fixed in formaldehyde 3.7% v/v and analyzed by light microscopy as described below.

### 2.3. Lipid droplet staining

Fixed parasites were centrifuged on glass slides at 30xg for 5 minutes. Slides were then washed with distilled water and subsequently kept in a 60% isopropanol solution for 5 minutes. Next, the slides were immersed in Oil Red O solution for 5 minutes. The slides were washed with distilled water and subsequently were mounted in aqueous medium, and the LDs marked by Oil Red O were quantified in 50 cells per slide using optical microscopy (Araújo-Santos et al., 2014).

### 2.4 Parasitic viability

Following treatment with PUFAs, *Leishmania* promastigotes were placed on 96-well flat-bottom plates in the presence of tetrazole salt (XTT) (ROCHE Applied Science) and incubated at 26°C for 4 hours in the dark. Next, XTT reduction by mitochondrial metabolism was evaluated by quantifying optical density on a plate reader (spectrophotometer) (Varioskan, ThermoScientific) (Fig. S1).

### 2.5 Lipid extraction to identify PUFAs and eicosanoids in parasite cell extract

After stimulation with AA, EPA or DHA, parasites isolated by centrifugation were subjected to hypotonic lysis in a 1:1 solution of deionized water and methanol at 4ºC. Culture supernatants were diluted at the same volume ratio in methanol at 4°C. Samples were then stored at −80ºC and sent for eicosanoid quantification at CEQIL, the Center of Excellence in Lipid Quantification and Identification (FCFRP-USP), using liquid chromatography/mass spectrometry (LC/MS) on a Nexera-TripleTOF® 5600+ Target Liquid Chromatography Tandem Mass Spectrometry (LC-MS/MS) system (SCIEX, Foster City, California). Next, oxylipid extraction was performed using the SPE (Solid Phase Extraction) method according to a previously described protocol (Sorgi et al., 2018). After lipid extraction, specimens were transferred to autosampler vials, and 10 μL of each sample were injected into the LC-MS/MS system as previously described by Sorgi et al., 2018. Final concentrations of oxylipids in culture extract and supernatants were quantified in accordance with standardized parameters (Sorgi et al., 2018).

### 2.7 In silico identification of enzymes involved in Leishmania eicosanoid metabolism

At least two enzymes related to the production of lipid mediators have been identified in Leishmania: GP63 (human-like COX) (Estrada-Figueroa et al., 2018) and prostaglandin F2α synthase (PGFS) (Araújo-Santos et al., 2014; Kabututu et al., 2003; Tavares et al., 2021). Initially, we performed a search for annotated nucleotide sequences characteristic of GP63 and PGFS using *L. major* as a reference species in GenBank. Using the BLASTn tool, complete genome sequences were identified and selected in the following species: *L. donovani, L. infantum, L. amazonensis, L. braziliensis, L. panamensis, L. mexicana, L. major* and *T. cruzi*. Only coding sequences (CDS) were considered for analysis (Table S1, S2). Multiple alignment of the obtained protein sequences was performed using the Clustal W (Codons) method. The Molecular Evolutionary Genetics Analysis (MEGA X) program was employed to construct a phylogenetic tree via the Unweighted Pair-Group Method with Arithmetic Mean (UPGMA) using 1000 bootstraps (Kumar et al., 2018).

### 2.8 3D modeling of GP63 and PGFS proteins in Old and New World *Leishmania* species

The Protein Data Bank (PDB) was used to search for *L. major* crystallographic structures in order to identify the structures of the GP63 and PGFS proteins (Berman, 2000); PDB IDs: 1LML for GP63 (Schlagenhauf et al., 1998) e PDB IDs: 4F40 for PFGS (Moen et al., 2015). The prediction of protein structures was then performed in other *Leishmania* spp using the Interactive Threading Assembly Refinement (I-TASSER) bioinformatics method (Zhang, 2015). Using these structures, the PyMOL program was employed to create overlapping models and analyze the active site residues of these enzymes (Yuan et al., 2017).

### 2.9 Western blot

Parasites in either stationary or logarithmic growth stages were lysed at a concentration of 2×10^8^/mL in RIPA solution. Next, the total amount of protein was quantified using the Pierce BCA protein assay (Thermo Scientific). Total proteins were separated by electrophoresis on a 10% polyacrylamide SDS gel and then transferred to nitrocellulose membranes. Membranes were blocked in Tris saline buffer (TBS) containing 0.1% Tween 20 (TT) plus 5% milk for 1 h, followed by incubation with *L. infantum* anti-PGFS (1:1000) overnight (Araújo-Santos et al., 2014). The primary antibody was then removed, and the membranes were washed five times in TT followed by incubation with the secondary antibody (goat anti-mouse) (SeraCare’s KPL Catalog 074-1806) conjugated to peroxidase (1:5000) for 1h. Finally, the membranes were washed 5x again and then incubated with Western Blotting substrate (Thermo Scientific Pierce ECL, Amersham, UK).

### 2.10 GP63 immunoassays

Protein extracts of *Leishmania* in logarithmic and stationary phase were submitted to ELISA to measure GP63 production. Briefly, 96-well immunoassay plates were sensitized with 30µg of *Leishmania* protein overnight at 4°C. Then, nonspecific binding was blocked with 0.1% PBS Tween 20 (PBS-T) plus 5% milk for 2 hours. After blocking, the plates were incubated with anti-GP63 (1:50) (Catalog # MA1-81830 Thermofisher) and incubated overnight at 4ºC. Next, the plates were washed with PBS-T and incubated with the secondary antibody (SeraCare’s KPL Catalog 074-1806) conjugated with peroxidase (1:2000) for 1 hour at room temperature. Finally, the plates were incubated with 3,3’5,5’-Tetramethylbenzidine (TMB) for 30 minutes, after which the reaction was stopped using 3M HCl. Plates were read on a microplate reader (Molecular Devices Spectra Max 340PC) a wavelength of 450nm.

### 2.11 Statistical analysis

Statistical analyses were performed using GraphPad-Prism v8.0 software (GraphPad Software, San Diego, CA-USA). All obtained data are represented as means ± standard error of the mean. Statistical analysis was performed using ANOVA or the Student Newman-Keuls test when comparing two groups, while Kruskal-Wallis was employed for three or more groups, at a significance level of p < 0.05. All experiments were performed in triplicate.

## 3. Results

### 3.1 Lipid droplets differ in quantity among *Leishmania* species

Our previous work demonstrated increasing numbers of LDs as *L. infantum* parasites differentiated into metacyclic forms in vitro (Araújo-Santos et al., 2014). Here, we also found higher numbers of LDs during the differentiation process in axenic cultures of *L. braziliensis* and *L. infantum*, but not in *L. amazonensis* (Fig. 1).

**Figure 1.**
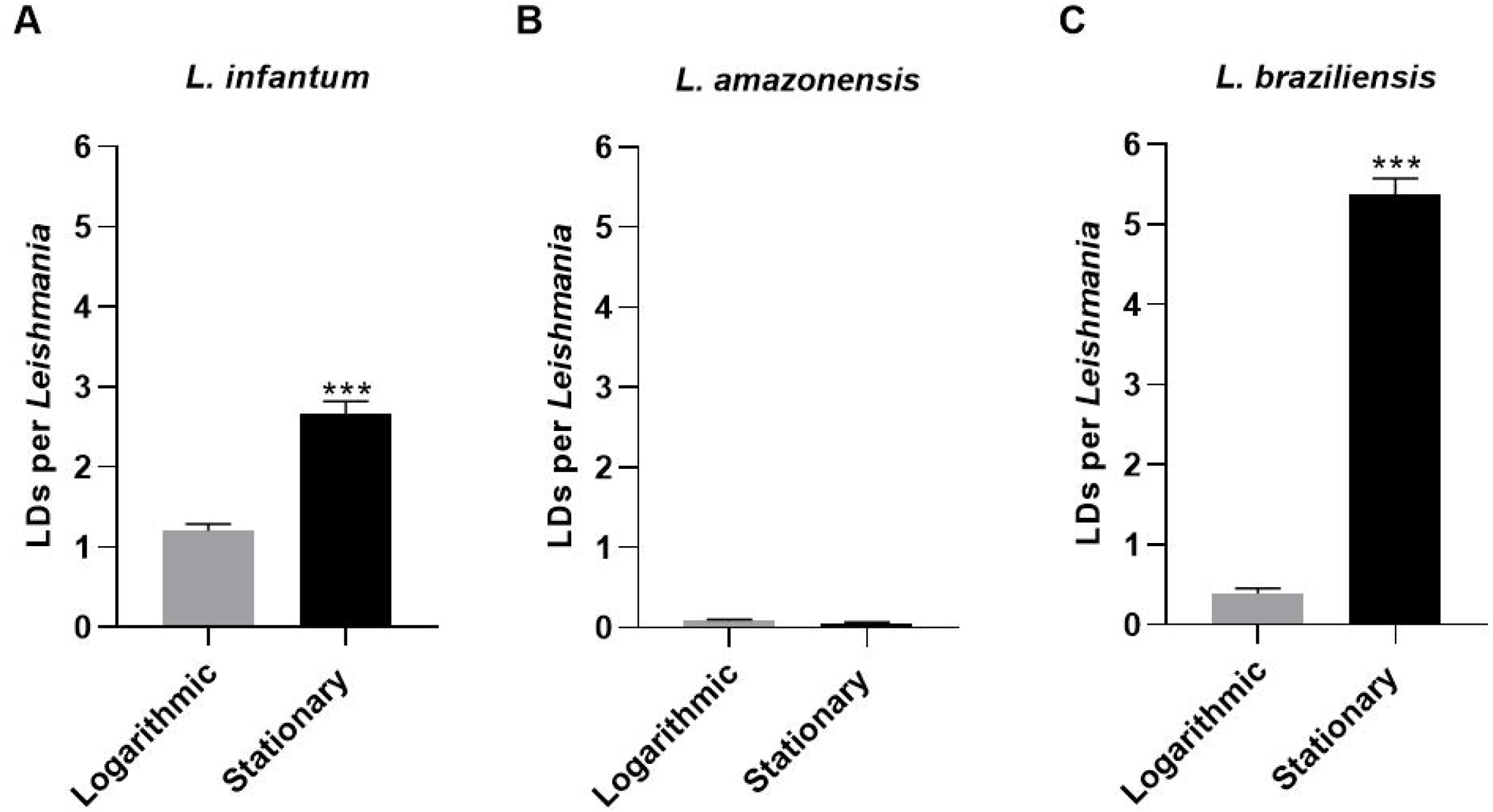
Lipid droplets in procyclic forms of *Leishmania spp*. Parasites in logarithmic and stationary growth phases were labeled with Oil Red O to count LDs. Data shown represent the mean ± standard error of LDs in (A) *L. infantum*, (B) *L. amazonensis*, and (C) *L. braziliensis*. ***, p<0.0001 using student Mann-Whitney test for multiple comparison by pairs.

### 3.2 PUFA stimulation increases lipid droplet formation in some *Leishmania* spp

LD formation was affected by stimulation with PUFAs; AA induced the formation of LDs in *L. braziliensis* and *L. infantum*, but not in *L. amazonensis* (Fig. 2). LD formation per parasite was found to be dose-dependent with regard to stimulation with EPA and DHA, as a statistically significant linear trend was observed for both *L. braziliensis* and *L. infantum*; however, no effect on LD formation was observed in *L. amazonensis* (Fig. 2 and Fig. S2).

**Figure 2.**
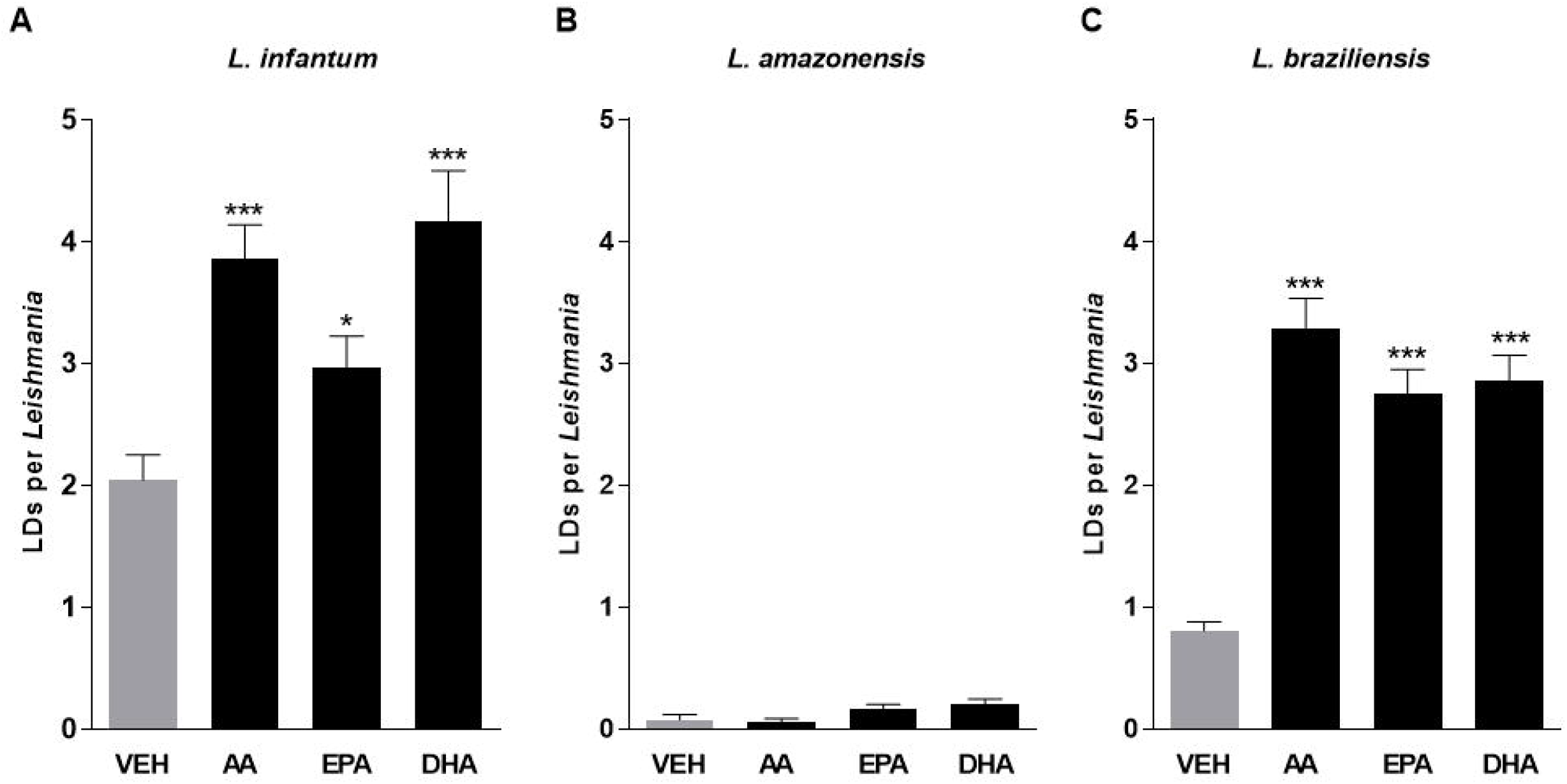
Polyunsaturated fatty acids increase the formation of lipid droplets in procyclic forms of *Leishmania*. Logarithmic growth phase promastigotes of (A) *L. infantum* (B) *L. amazonensis* and (C) *L. braziliensis* were stimulated with ethanol (vehicle) or AA (15 µM), EPA (30 µM) or DHA (30 µM) for 1 hour, and then stained with Oil Red O to quantify LDs. Bars represent means ± SEM of LDs per parasite. *** and * represent p<0.0001 and p<0.05, respectively, for multiple pairwise comparisons between stimuli and the vehicle using the Student-Newman-Keuls test. AA: Arachidonic acid; EPA: Eicosapentaenoic acid; DHA: Docosahexaenoic acid.

### 3.3 Comparison of eicosanoid metabolism enzymes in Old and New World *Leishmania spp*

Proteins associated with eicosanoid metabolism are present in *Leishmania spp*., such as a COX-like enzyme, previously known as GP63, and PGFS (Tavares et al., 2021). We performed a comparison of the primary protein structure sequences in silico (Fig. S3). In addition, the tertiary structures of these reference proteins exhibited high structural similarities across *Leishmania spp*, as well as *T. cruzi* (Figs S4, S5). Alignment and phylogenetic analysis of GP63 and PGFS indicated notable similarity and homology between the Old and New World *Leishmania* spp analyzed with respect to clinical manifestations of disease (viscerotropic, dermotropic or mucotropic; Fig. S3; Figs 3A, B). In addition, nonsynonymous mutations were identified in the amino acid residues at the active sites of GP63 (PDB ID: 1LML) and PGFS (PDB ID: 4F40). *Leishmania infantum* and *L. donovani* shared mutations in GP63 protein residues. The dermotropic species presented same residues in the active site of the GP63 enzyme, while the species with mucosal tropism presented exclusive mutations in the active site of GP63 (Fig. 3C). Furthermore, most of the PGFS residues among viscerotropic, dermotropic and mucotropic species were conserved, except for the K204 residue that was altered in *L. braziliensis* and *L. panamensis*, both mucotropic species (Fig. 3D).

**Figure 3.**
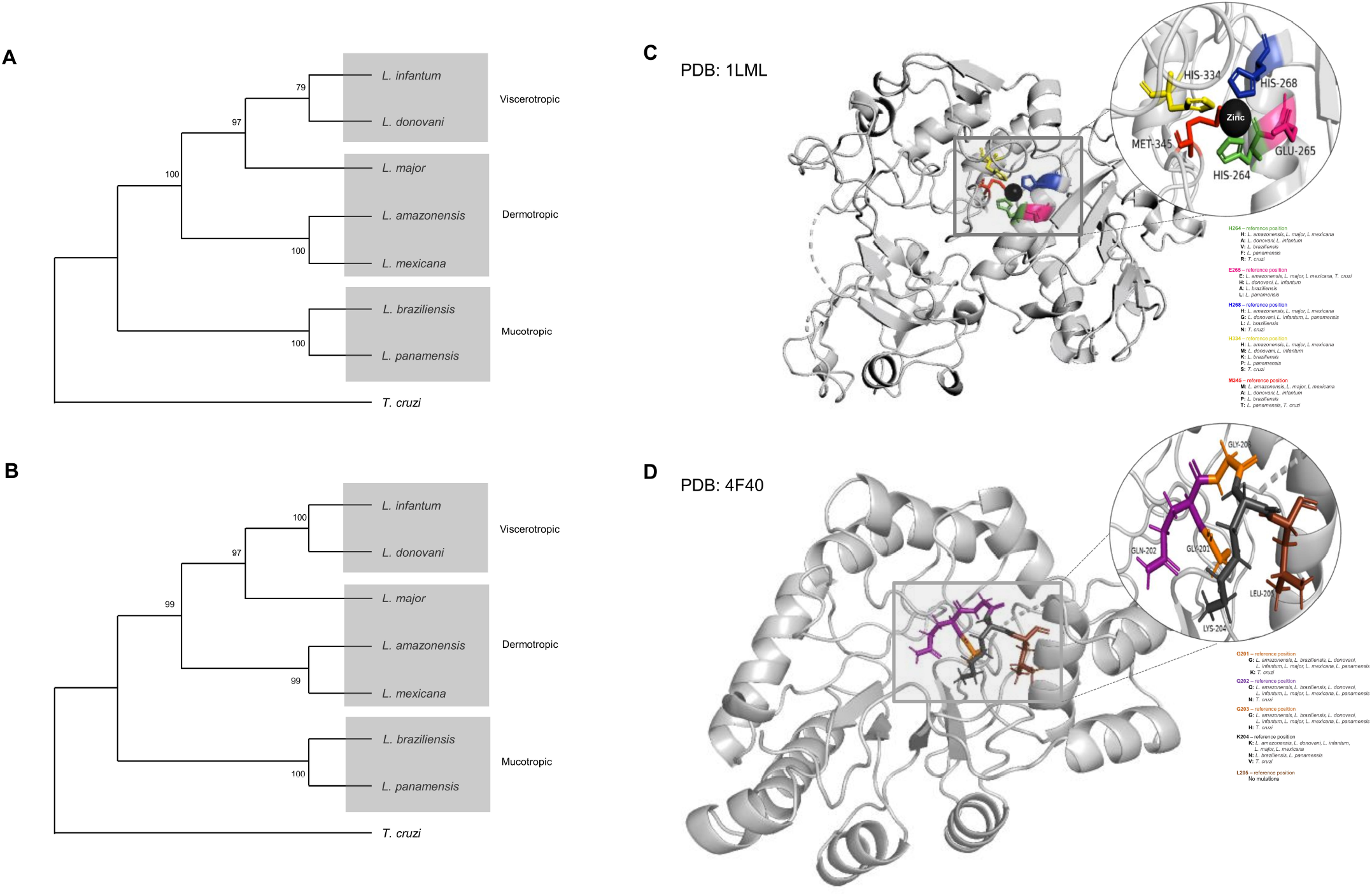
Homology of GP63 and PGFS proteins across different *Leishmania spp*. GP63 and PGFS genes were identified in the reference genomes of etiologic pathogens related to human leishmaniasis. Phylogenetic trees of (A) GP63 and (B) PGFS were constructed using the UPGMA method (MEGA X software), considering 1000 bootstraps. Residues at the active sites of (C) GP63 and (D) PGFS proteins are highlighted by colors depicted in three-dimensional structures. Non-synonymous mutations among *Leishmania spp*. are listed.

We then compared the gene expression of these two enzymes involved in the production of eicosanoids in different *Leishmania* species. GP63 was more produced in log-stage *L. infantum* and *L. braziliensis* parasites, but not in *L. amazonensis* (Fig. 4A-C). In addition, PGFS was more produced in stationary parasites compared to log-stage procyclic forms in all three species of *Leishmania* (Fig. 4D-G).

**Figure 4.**
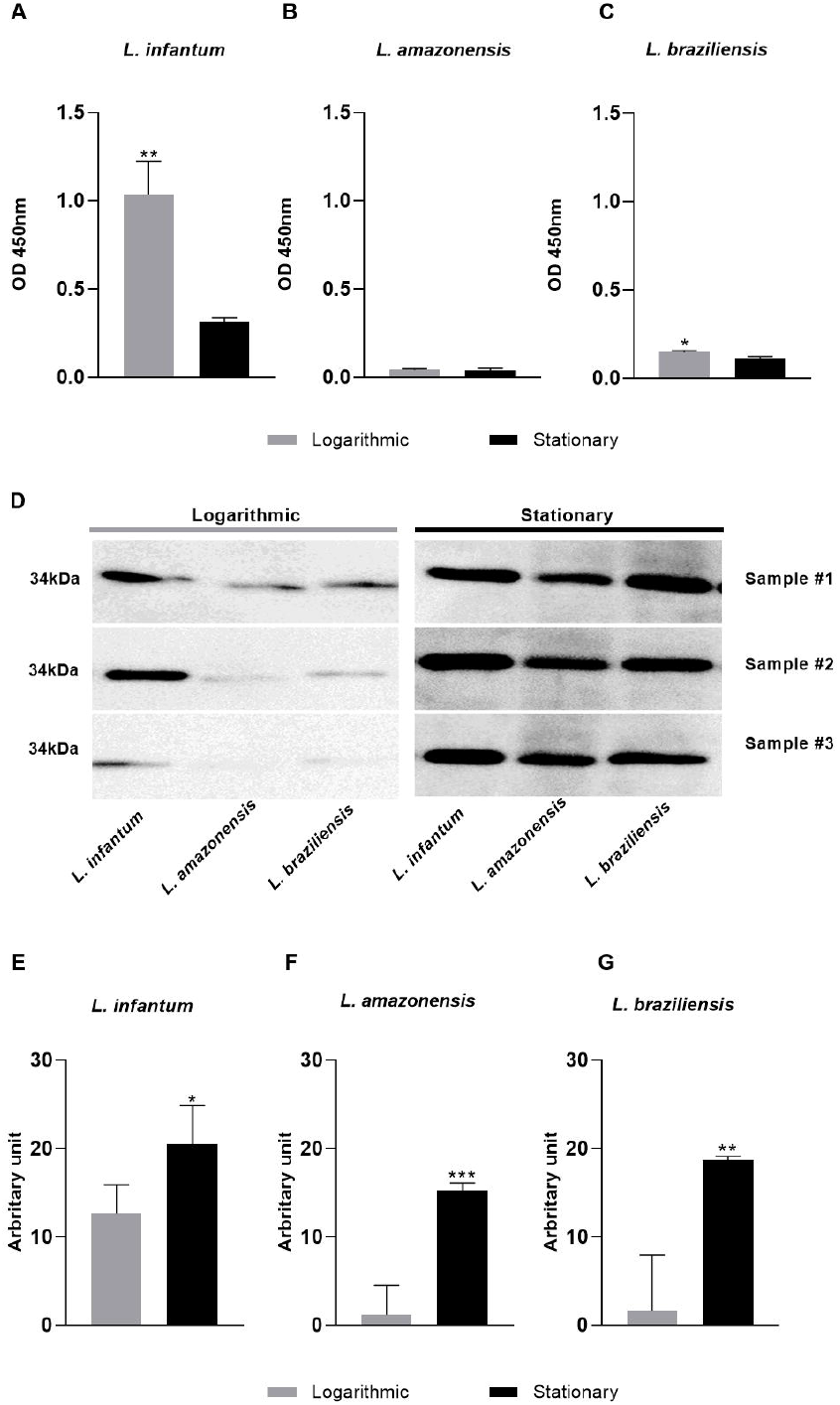
GP63 and PGFS protein production in logarithmic and stationary axenic stages of *Leishmania spp*. Parasites in logarithmic and stationary growth phases were lysed to measure GP63 protein production in (A) *L. infantum*, (B) *L. amazonensis*, (C) *L. braziliensis* by immunoassay. Total protein (30µg) was incubated with anti-PGFS (1:500) for Western blot analysis. (D) Immunoblot comparing the abundance of PGFS in logarithmic and stationary stages incubated with anti-PGFS (1:1000) in (E) *L. infantum*, (F) *L. amazonensis*, (G) *L. braziliensis*. * p<0.05 for multiple pairwise comparisons between stimuli and the vehicle using the Student-Newman-Keuls test.

### 3.4 Qualitative effects of PUFAs on eicosanoid production in *Leishmania spp*

Trypanosomatids possess enzymatic machinery for the metabolization of PUFAs to specialized and conventional lipid mediators (Colas, 2018; Paloque et al., 2019b; Tavares et al., 2021). Herein we used LC/MS to evaluate the presence of lipid mediator precursors, as well as eicosanoids, in *L. infantum, L. amazonensis* and *L. braziliensis* treated or not with AA, EPA or DHA. In all, 41 lipid mediators were analyzed in cell extracts and axenic culture supernatants. The presence of twelve bioactive lipids was identified: 15-keto-PGE_2_, LXA_4_, PGD_2_, PGE_2_, 5-HETE, AA, 12-HETE, 8-HETE, 11-HETE, EPA, 15-HETE, PGF_2α_ (Table 1 and Fig. 5). However, mediators LTC_4_, PGB_2_, 6-keto-PGF1a, 17-RvD1, 12-oxo-LTB_4_, 20-OH-PGE_2_, TXB_2_, RvD1, RvD2, RvD3, LTB_4_, LTD_4_, LTE_4_, 6-trans-LTB_4_, 11-trans-LTD_4_, PDx, Maresin, PGJ_2_/PGA_2_, RvE1, 15-deoxy-PGJ_2_, 5-oxo-ETE, 20-HETE, 5,6-DiHETE, 12-oxo-ETE, 15-oxo-ETE, 11,12-DiHETrE, 14,15-DiHETrE, 5,6-DiHETrE, 20-OH-LTB4 were not identified. As expected, AA-derived eicosanoids were the most prevalent in *Leishmania* extracts (Fig. 5).

**Table 1.** Quantitation of eicosanoids and their precursors in cell extract and supernatant of *Leishmania spp*. by LC-MS.

| Lipids | Standards* | Mass (m/z) |  | Lipid concentrations in cell extract and supernatant by *Leishmania* spp (ng/mL)* |  |  |  |  |  |
| --- | --- | --- | --- | --- | --- | --- | --- | --- | --- |
|  |  | Precursor ion (m/z) | Fragment ion (m/z) | *L. infantum* |  | *L. amazonensis* |  | *L. braziliensis* |  |
|  |  |  |  | C | S | C | S | C | S |
| **PGF<sub>2α</sub>** | PGF<sub>2α</sub>-d4 | 353 | 309.2179 | 0.17 | 0.27 | <0.01 | 4.68 | <0.01 | 1.62 |
| **15-Keto-PGE<sub>2</sub>** | PGE<sub>2</sub>-d4 | 349 | 287.2017 | 2.82 | 43.80 | 1.32 | 36.12 | 1.12 | 33.89 |
| **PGE<sub>2</sub>** | PGE<sub>2</sub>-d4 | 351 | 189.1285 | 2.30 | 11.14 | 1.46 | 34.19 | 6.31 | 77.74 |
| **PGD<sub>2</sub>** | PGD<sub>2</sub>-d4 | 351 | 189.1285 | 1.70 | 13.29 | 2.69 | 8.56 | 1.09 | 15.21 |
| **LXA<sub>4</sub>** | LXA<sub>4</sub>-d5 | 351 | 217.1598 | 7.27 | 17.59 | 10.46 | 35.05 | 11.66 | 22.76 |
| **15-HETE** | 15-HETE-d8 | 319 | 175.1492 | 6.20 | 50.66 | 16.23 | 187.49 | 16.34 | 182.75 |
| **12-HETE** | 12-HETE-d8 | 319 | 179.1078 | 3.14 | 26.65 | 8.26 | 47.16 | 7.57 | 30.52 |
| **11-HETE** | 12-HETE-d8 | 319 | 167.1084 | 4.52 | 25.52 | 9.93 | 65.03 | 11.59 | 49.60 |
| **8-HETE** | 12-HETE-d8 | 319 | 155.0714 | 3.82 | 22.01 | 9.00 | 55.83 | 9.88 | 32.00 |
| **5-HETE** | 5-HETE-d8 | 319 | 115.0401 | <0.01 | 58.94 | 18.74 | 81.46 | 20.79 | 75.68 |
| **EPA** | AA-d11 | 301 | 257.2275 | 1.48 | 14.54 | 14.54 | 43.07 | 3.99 | 8.10 |
| **AA** | AA-d11 | 303 | 259.2447 | 41.59 | 365.81 | 59.94 | 419.78 | 57.27 | 252.77 |

AA: Arachidonic Acid; EPA: Eicosapentaenoic acid; LX: Lipoxin; PG: prostaglandin; HETE: Hydroxyeicosatetraenoate; C: Cell extract; S: Supernatant;
\* Standard molecules containing Deuterium (d) atoms was used as an internal standard for the quantification of lipids by LC-mass spectrometry (MS).

**Figure 5.**
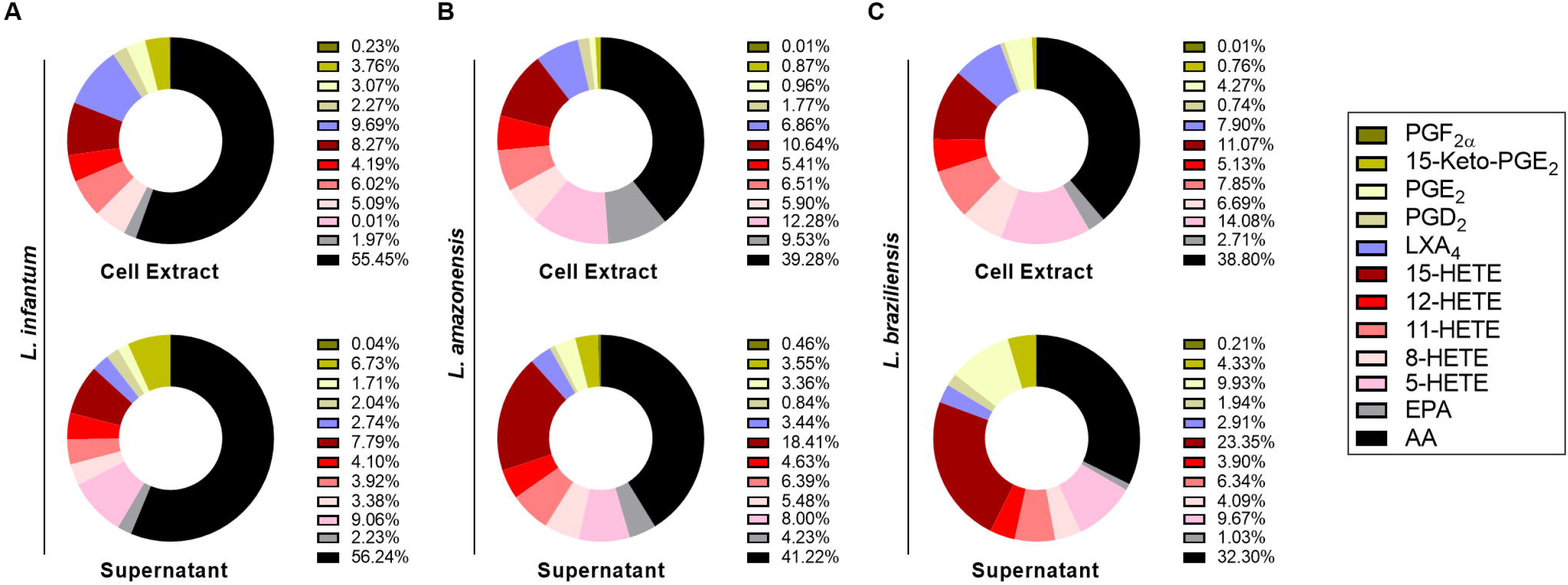
Identification of lipid mediators in different *Leishmania* spp. Average distribution of the 10 abundant eicosanoids in the cell extract and supernatant of procyclic promastigotes of (A) *L. infantum*, (B) *L. amazonensis* and (C) *L. braziliensis* in logarithmic growth phase (with cell types labeled above each pie chart) stimulated with AA for 1 hour, displayed as parts of the whole. Data are in percentage of ηg/mL of total lipid detected by LC-MS. EPA: Eicosapentanoic acid; 15-HETE: 15-hydroxyeicosatetraenoic acid; 8-HETE: 8-hydroxyeicosatetraenoic acid; 11-HETE: 11-Hydroxyeicosatetraenoic acid; 12-HETE: 12-hydroxyeicosatetraenoic acid; AA: Arachidonic acid; PGF_2α_: Prostaglandin F_2α_; 15-keto-PGE_2_: 15-keto-Prostaglandin E_2_; LXA_4_: Lipoxin A_4_; PGD_2_: Prostaglandin D_2_; 5-HETE: 5-hydroxyeicosatetraenoic acid.

## 4. Discussion

While lipid mediators and LDs have a potential role in the pathogenicity of *Leishmania* infection, the literature contains scarce data on eicosanoid metabolism in these parasites. Here we compared eicosanoid metabolism and LD formation in response to PUFAs in different species of *Leishmania* associated with distinct clinical forms. Our data show that, in contrast to *L. amazonensis*, LD formation can be modulated by PUFA stimulation in *L. infantum* and *L. braziliensis*. However, with regard to eicosanoid production, the same eicosanoids were detected across all three New World species when stimulated with AA, yet *L. amazonensis* did not respond similarly to the other two species investigated under stimulation with EPA and DHA. The mechanisms of regulation of the production of LDs in *Leishmania* are still poorly understood. *Trypanosomatids* have only two proteins related to the LD formation, the lipid droplet kinase (LDK) and *Trypanosoma brucei* Lipin (TbLpn) (Dawoody Nejad et al., 2018; Flaspohler et al., 2010; Tavares et al., 2021). More studies will be necessary to investigate the presence and function of both LDK and TbLpn in *Leishmania* species. We can hypothesize that decrease in the activity in *L. amanozensis* could explain the inability of this parasite species to respond to PUFAs stimuli with the formation of LDs.

LDs are organelles that synthesize lipid mediators in a variety of cell types (Bozza et al., 2011). However, the role of LDs in eicosanoid production in parasites remains poorly understood (Tavares et al., 2021). In *L. infantum*, LDs are sites of production for PGF_2α_ (Araújo-Santos et al., 2014), but the literature contains scare reports on the presence of enzymes related to the lipid metabolism of eicosanoids or bioactive lipids in other protozoa. Herein, we found that AA modulated the formation of LDs, which suggests the accumulation of AA in LDs that may serve as a platform for the synthesis of eicosanoids in some protozoa of the genus *Leishmania*.

Regarding the formation of lipid mediators, it is known that enzymatic machinery in protozoa is responsible for the synthesis of these bioactive compounds, such as GP63 (Estrada-Figueroa et al., 2018) and PGFS (Alves-Ferreira et al., 2020; Araújo-Santos et al., 2014; Kabututu et al., 2003; Tavares et al., 2021; Toledo et al., 2016). However, the enzymes that convert AA into lipid mediators have not been adequately studied. In *L. mexicana*, a GP63 protein was shown to be analogous to COX-2 (Estrada-Figueroa et al., 2018). These protein metalloproteases are described as the main surface antigen expressed in promastigotes of different *Leishmania* species (Isnard et al., 2012). Although the genes encoding the GP63 metalloproteases are organized in tandem (Ivens et al., 2006; Peacock et al., 2007), it has not been demonstrated whether COX-2 activity would arise from all of these encoded proteins. The production of PGF2α, which enzyme PGFS is responsible for synthesizing (Kubata et al., 2007), activates the PGF2α receptor, triggering the COX pathway (Ueno and Fujimori, 2011). While little is known about the role of parasite-derived PGFS, some studies suggest the potential role of this eicosanoid in host-parasite interaction (Alves-Ferreira et al., 2020; Araújo-Santos et al., 2014). The analysis of the viscerotropic and dermotropic *Leishmania* species point out some genetic differences related to parasite tropism (Ait Maatallah et al., 2022). However, a relationship between the genetics of the parasites and the pathophysiology is still necessary. In this sense, identifying the differences in the production of eicosanoids can help to explain the distinct clinical manifestations caused between different species of leishmania, since the mechanisms of eicosanoid action during inflammation are well knowns (Bozza et al., 2011, 2009). Our comparison of protein sequences and GP63 and PGFS active site residues in both Old and New World *Leishmania* species revealed surprising similarity between these enzymes in accordance with the clinical form of disease caused by the parasite. Nonetheless, additional studies are needed to verify if, in fact, this similarity could be related to GP63 and PGFS production, and to the development of a polarized response according to the clinical manifestation. Indeed, some studies in humans have shown altered production of lipid mediators depending on the cutaneous or visceral form of disease (Araújo-Santos et al., 2014; Malta-Santos et al., 2020).

An important aspect of our evaluation focused on the production of enzymes involved in the metabolism of eicosanoids during the metacyclogenesis of different New World *Leishmania* species. The COX-like enzyme GP63 production reduces during *L. braziliensis* and *L. infantum* promastigotes differentiation, but no differences are observed in *L. amazonensis*. In the next AA metabolism step, PGFS production increases during parasite differentiation in different *Leishmania* species. This finding may indicate that PGFS could influence the virulence of infective forms of *Leishmania spp*. Considering the differences in the structure of and family of genes encoding these enzymes, further studies are needed to correlate these differences with the enzymatic activity exhibited by PGFS in *Leishmania*.

While some studies have advanced the understanding of lipid metabolism, the enzymes involved in parasites remain poorly described; however, knowledge on which metabolites are produced by zoonotic parasites is expanding (de Almeida et al., 2018; Estrada-Figueroa et al., 2018; Paloque et al., 2019b; Toledo et al., 2016). Here, we investigated the metabolites produced by different *Leishmania* species, observing increased production of lipid mediators of the HETE class. Importantly, little is known about the role played by these metabolites during the host-parasite interaction process. Additional studies may shed light on whether these should be considered virulence factors and thus may serve as intervention targets, in addition to whether their currently unidentified receptors could elucidate mechanisms of pathogenicity. The present findings serve to open perspectives by providing evidence on how PUFAs lead to the modulation of LD formation in different Old and New World *Leishmania* species. We believe that our qualitative overview of lipid mediators potentially produced by these parasites contributes to the base of knowledge surrounding antiparasitic drug development.

## Author Contributions

Conceived and designed the experiments: VMB, CAS, TA-S. Data collection: MVC, VST, RSOS, YMFSA, CAS, TA-S; Data analysis: MVC, VST, RSOS, CAS, JLB, VMB, TA-S; Contributed materials/analysis tools: LHF, CAS, JLB, VMB, TA-S; Wrote the paper: YMFSA and TA-S.

## Acknowledgments

This work was supported by the Brazilian National Council for Scientific and Technological Development (CNPq) (grant number #422696/2016-1) to TA-S and (grant number #431857/2018-0) to VMB; Financiadora de Estudos e Projetos (FINEP) (grant number #0418000600) to TA-S; Fundação do Amparo à Pesquisa do Estado de São Paulo (FAPESP) (EMU grant number #2015/00658-1) to LHF; Fundação de Amparo à Pesquisa do Estado da Bahia (FAPESB) to RSOS, YMFSA; and Coordenação de Aperfeiçoamento de Pessoal de Nível Superior (CAPES). The authors would like to thank Andris K. Walter for critical analysis, English language revision and manuscript copyediting assistance.

