## Supplementary material for "Polyunsaturated fatty acids alter the formation of lipid droplets and eicosanoid production in *Leishmania* promatigotes": Suplemmental material

### Supplementary table 1. GP63 nucleotide sequences used in the study

| Species | GenBank Code | region (begin - end) | Size (bp) | UniprotKB code |
|---|---|---|---|---|
| *Leishmania infantum* | FR796442.1 | 222401 - 224197 | 1800 | Q6LA77 |
| *Leishmania donovani* | CP029509.1 | 257944 - 259743 | 1800 | A0A3S7WR60 |
| *Leishmania major* | Y00647.1 | 199 - 2007 | 1809 | P08148 |
| *Leishmania amazonensis* | CP040138.1 | 179953 - 181761 | 1809 | No annotation |
| *Leishmania mexicana* | NC_018314.1 | 180317 - 182125 | 1809 | E9AN54 |
| *Leishmania braziliensis* | LS997609.1 | 224752 - 226554 | 1803 | A0A3P3YZR7 |
| *Leishmania panamensis* | AF037166.1 | 1 - 1770 | 1770 | O46312 |
| *Trypanosoma cruzi* | MKQG01002498.1 | 73636 - 75255 | 1630 | No annotation |

### Supplementary table 2. PGFS nucleotide sequences used in the study

| Species | GenBank Code | region (begin - end)* | Size (bp) | UniProtKB code |
|---|---|---|---|---|
| *Leishmania infantum* | FR796463.1 | 1034722 - 1035576 | 855 | A4I6Z4 |
| *Leishmania donovani* | FR799618.2 | 1062787 - 1063641 | 855 | E9BMZ2 |
| *Leishmania major* | FR796427.1 | 1050960 - 1051814 | 855 | P22045 |
| *Leishmania amazonensis* | CP040158.1 | 1024015 -1023161 | 855 | No annotation |
| *Leishmania mexicana* | FR799583.1 | 1026690 -1027544 | 855 | E9B215 |
| *Leishmania braziliensis* | FR799006.1 | 1114033 - 1114887 | 855 | A4HJJ7 |
| *Leishmania panamensis* | CP009400.1 | 936089 - 936943 | 855 | A0A088RXB1 |
| *Trypanosoma cruzi* | AAHK01000429.1 | 11770 - 12618 | 849 | Q4DJ07 |

*gene identified in the reverse complementary sequence

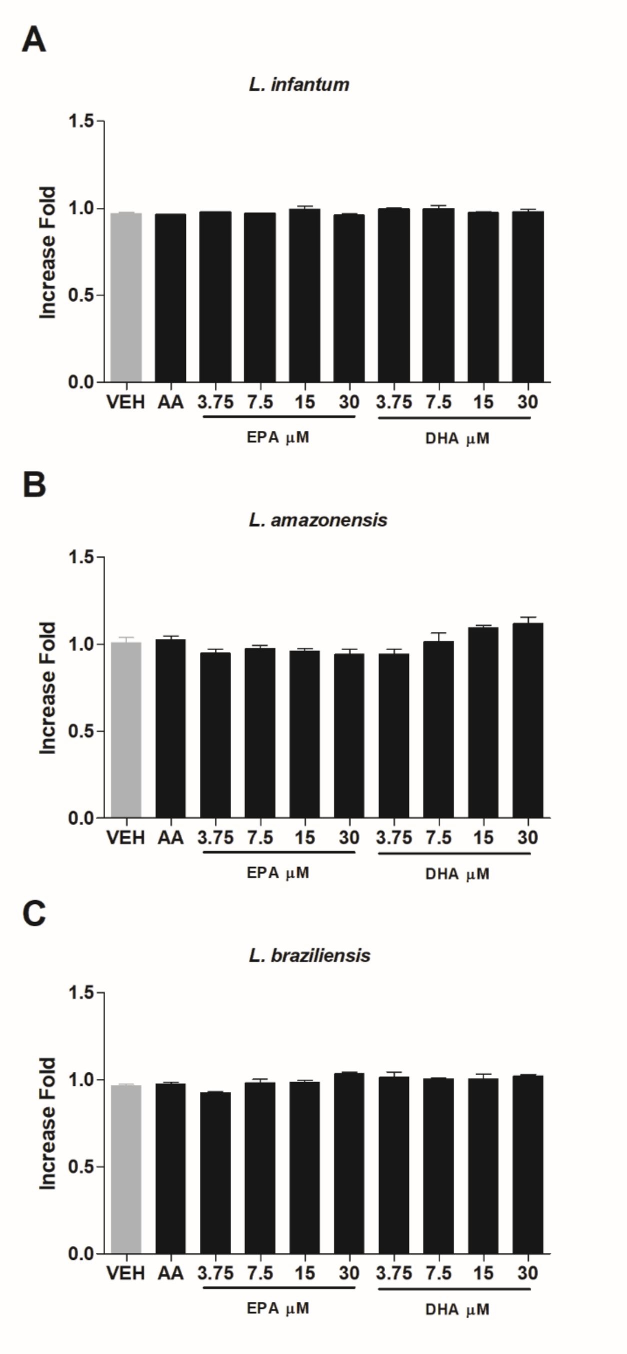


**Supplementary figure 1. Polyunsaturated fatty acids stimulation does not affect *Leishmania* viability.** Procyclic promastigotes of (A) *L. infantum*, (B) *L. amazonensis* and (C) *L. brasiliensis* in logarithmic growth phase were stimulated with AA, EPA or DHA for 1 hour. Next, tetrazolium salt (XTT) reduction was measured by spectrophotometry. Data are represented as means ± standard error of optical density readings.

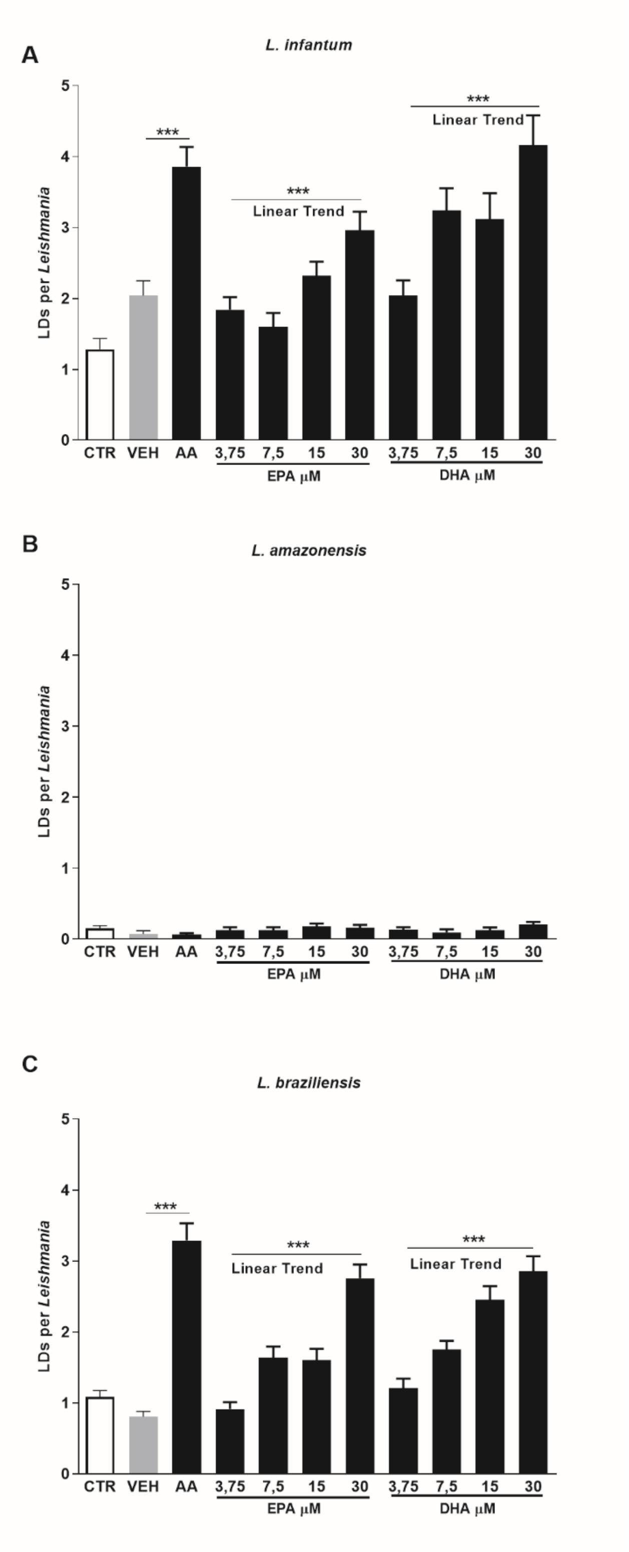


**Figure 2. Polyunsaturated fatty acids increase the formation of lipid droplets in procylic forms of *Leishmania*.** Logarithmic growth phase promastigotes of (A) *L. infantum* (B) *L.*

*amazonensis* and (C) *L. braziliensis* were stimulated with ethanol (vehicle) or AA (15 µM), EPA (3.75, 7.5, 15 or 30 µM) or DHA (3.75, 7.5, 15 or 30 µM) for 1 hour, and then stained with Oil Red O to quantify LDs. Bars represent means ± SEM of LDs per parasite. *** represent $p<0.0001$, for pairwise comparison between AA and the vehicle using the Student´s t-test. The significance was tested by 1-way ANOVA with post-test linear trend to dose response stimuli. AA: Arachidonic acid; EPA: Eicosapentaenoic acid; DHA: Docosahexaenoic acid.

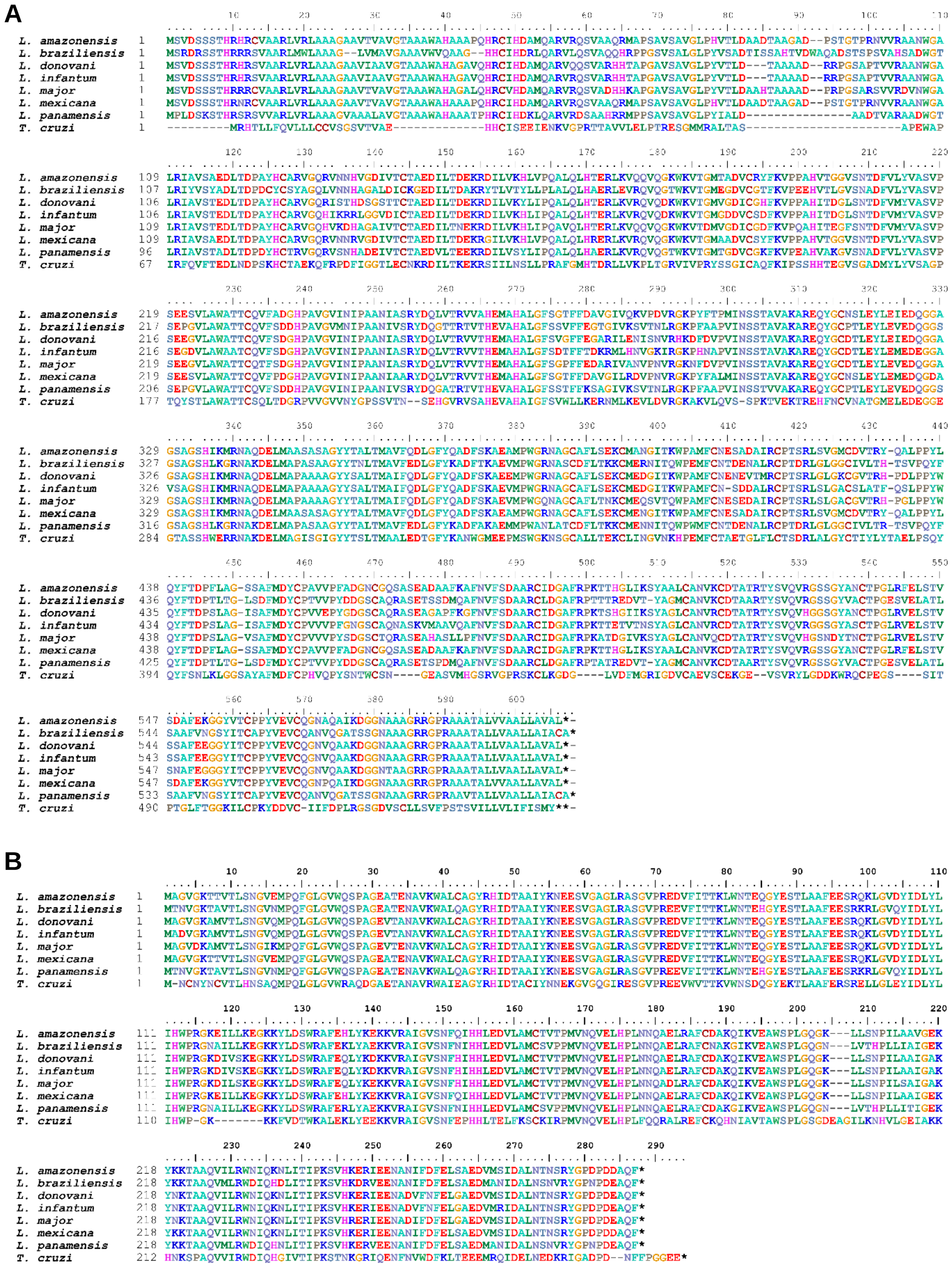


**Supplementary figure 3. Comparative analysis of the primary structure of GP63 and PGFS proteins across New and Old World *Leishmania* spp.** Protein sequences of (A) GP63 or (B) PGFS were aligned using the ClustalW algorithm.

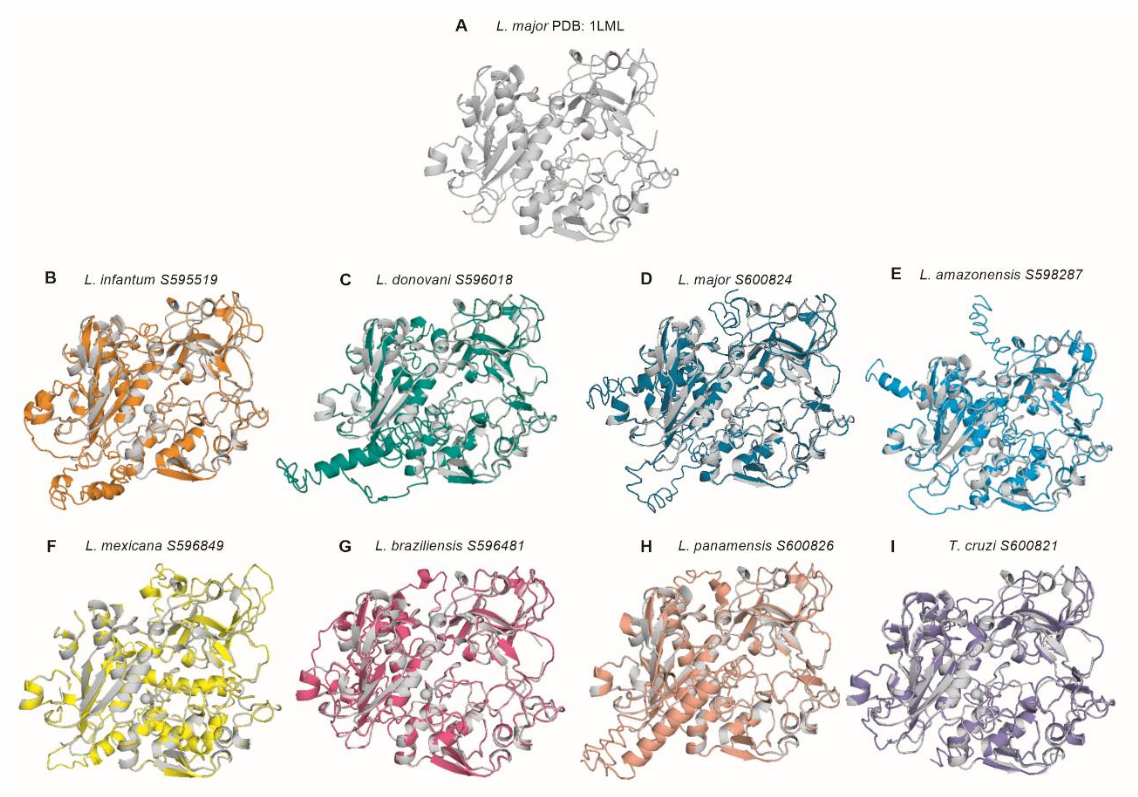


**Supplementary figure 4. Comparative analysis of the tertiary structure of GP63 protein in *Leishmania spp.* and *Trypanosoma cruzi.*** The GP63 protein was modeled using the I-TASSER algorithm and structures were aligned over the *L. major* protein (grey). GP63 tertiary structure overlap shows similarities between (A) *L. major* PDB: 1LML and (B) *L. infantum*, (C) *L. donovani*, (D) *L. major*, (E) *L. amazonensis*, (F) *L. mexicana*, (G) *L. braziliensis*, (H) *L. panamensis* and (I) *T. cruzi*.

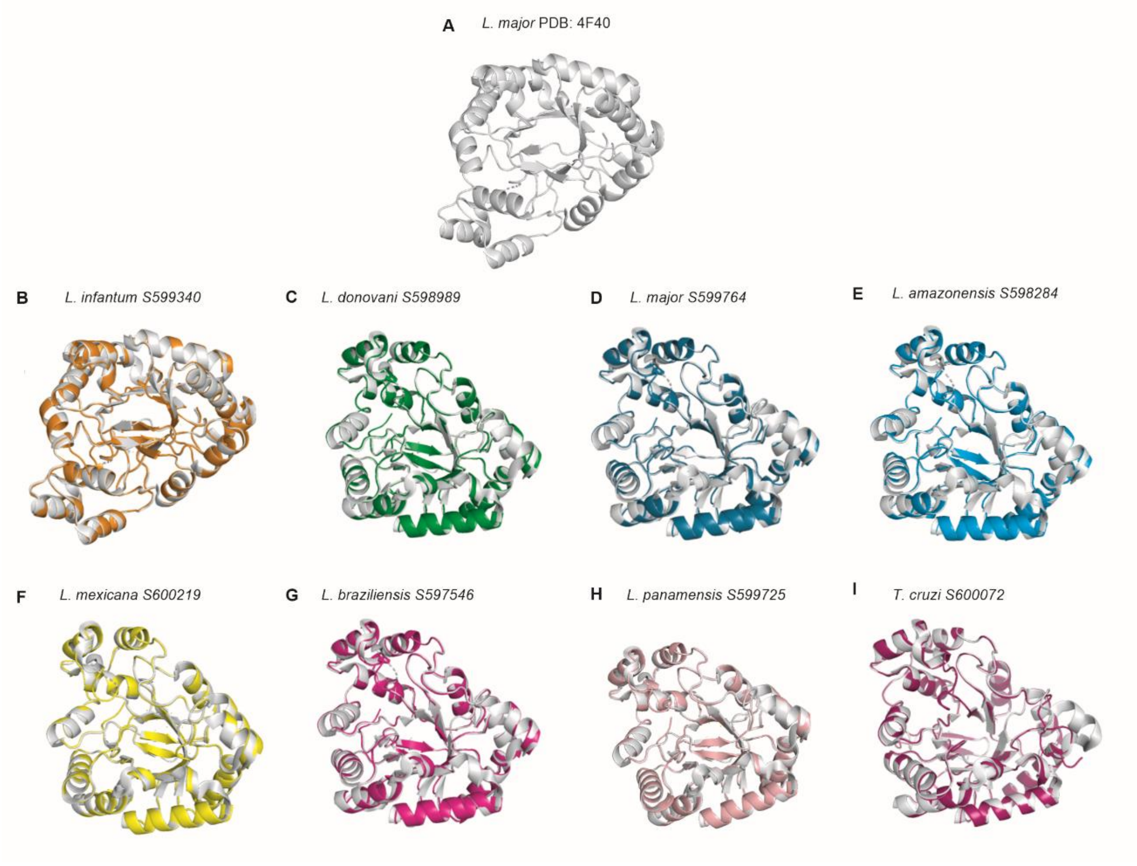


**Supplementary figure 5. Comparative analysis of the tertiary structure of PGFS protein in *Leishmania spp.* and *Trypanosoma cruzi.*** The PGFS protein was modeled using the I-TASSER algorism and structures were aligned over the *L. major* protein (grey). PGFS tertiary structure overlap shows similarities between (A) *L. major* PDB: 4F40 and (B) *L. infantum*, (C) *L. donovani*, (D) *L. major*, (E) *L. amazonensis*, (F) *L. mexicana*, (G) *L. braziliensis*, (H) *L. panamensis* and (I) *T. cruzi*.
